# MassVis*ion*: An open-source end-to-end platform for AI-driven mass spectrometry image analysis

**DOI:** 10.1101/2025.01.29.635489

**Authors:** Amoon Jamzad, Ayesha Syeda, Jade Warren, Martin Kaufmann, Natasha Iaboni, Christopher J. B. Nicol, John Rudan, Kevin Y. M. Ren, David Hurlbut, Sonal Varma, Gabor Fichtinger, Parvin Mousavi

## Abstract

Mass spectrometry imaging (MSI) combines spatial and spectral data to reveal detailed molecular compositions within biological samples. Despite their immense potential, MSI workflows are hindered by the complexity and high dimensionality of the data, making their analysis computationally intensive and often requiring expertise in coding. Existing tools frequently lack the integration needed for seamless, scalable, and end-to-end workflows, forcing researchers to rely on local solutions or multiple platforms, hindering efficiency and accessibility. We introduce MassVision, a comprehensive software platform for MSI analysis. Built on the 3D Slicer ecosystem, MassVision integrates MSI-specific functionalities while addressing general user requirements for accessibility and usability. Its intuitive interface lowers barriers for researchers with varying levels of computational expertise, while its scalability supports high-throughput studies and multi-slide datasets. Key functionalities include visualization, co-localization, dataset curation, dataset merging, spectral and spatial preprocessing, AI model training, and AI deployment on full MSI data. We detail the workflow and functionalities of MassVision and demonstrate its effectiveness through different experimental use cases such as exploratory data analysis, ion identification, and tissue-type classification, on in-house and publicly available data from different MSI modalities. These use cases underscore the MassVision’s ability to seamlessly integrate MSI data handling steps into a single platform, and highlight its potential to reveal new insights and structures when examining biological samples. By combining cutting-edge functionality with user-centric design, MassVision addresses longstanding challenges in MSI data analysis and provides a robust tool for advancing the user’s ability to achieve biologically-meaningful insights from MSI data.

## Introduction

Mass spectrometry imaging (MSI) is a technique that combines spatial and mass spectral data to facilitate detailed insights into the molecular composition of tissue sections. By mapping the spatial distribution of metabolites, lipids, and proteins, MSI has revolutionized applications in metabolomics, biology, and clinical diagnostics.^1^ In cancer research, MSI has proven invaluable for identifying molecular signatures of tumor tissues, as well in drug development and treatment monitoring.^2^ Its ability to detect molecular signatures without relying on external molecular markers like antibodies or dyes highlights its potential as a point-of-care histopathology tool for rapid tissue characterization. ^3^ Recent advancements, including emerging computational methods and improved analytical workflows, have expanded MSI’s utility in studying disease mechanisms and therapeutic responses, underscoring its growing role in both research and clinical applications.^4,5^

Artificial Intelligence (AI) is transforming data analysis across a wide range of scientific fields, including MSI.^6–9^ While traditional statistical approaches have been fundamental to MSI, the field is increasingly shifting toward AI-driven methods to address the complexity, high dimensionality, and large-scale nature of MSI data. AI algorithms may be broadly divided into two main categories: Unsupervised AI techniques are employed to reduce spectral dimensionality, facilitating visualization, exploration, and ion clustering;^10^ In contrast, supervised AI models are designed to predict tissue types based on known molecular signatures.^11^ Regardless of the algorithm, training and validation of AI models relies on well-curated MSI datasets, where spectra serve as data samples, and tissue labels are typically derived from a gold-standard modality, such as histopathology.^4,11^ While AI methods have proven effective for MSI analysis, their implementation often requires technical programming skills, posing a significant barrier for researchers without coding expertise. Many research groups develop custom in-house platforms to address their specific research needs, but challenges with reproducibility, accessibility, and standardization hinder their broader applicability in other settings.

Several commercial and open-source software platforms have been developed to facilitate and standardize the broader adoption of analytical methods in MSI analysis.^12–15^ Some provide advanced statistical and AI algorithms for downstream spectral analysis and ignore spatial information,^14^ while others focus exclusively on dataset curation, offering visualization and region-of-interest (ROI) selection functionalities. ^12^ However, these software platforms often fall short of supporting a complete end-to-end pipeline for AI integration, and as a result, researchers are compelled to rely on multiple software solutions for intermediate steps. This approach requires compatibility adjustments, slows down analysis, and increases the risk of errors, ultimately compromising pipeline efficiency and reliability. Furthermore, available software applications lack critical features such as flexible visualization to highlight local metabolomic variations, co-localization of MSI with histopathology images to guide ROI selection, and seamless merging of multiple datasets for multi-patient studies. Moreover, available platforms lack the capability to integrate trained AI models for prospective deployment, limiting the translation of experimental analysis to real-world clinical implementation. These shortcomings prevent researchers from fully leveraging the potential of MSI. Bridging these gaps requires a new generation of software capable of supporting end-to-end workflows, enabling seamless data integration and AI deployment, to maximize the impact and accessibility of MSI research.

Here, we introduce MassVision, an open-source comprehensive end-to-end solution for MSI analysis developed to address the above challenges. MassVision offers a wide range of functionalities, including data exploration and visualization, histopathology co-localization, dataset curation, multi-slide merging, spectral and spatial preprocessing, AI model training and evaluation, and deploying these models to the whole-slide MSI. Beyond its MSI-specific capabilities, MassVision is designed to be accessible, usable, flexible, and maintainable, ensuring seamless integration, long-term sustainability, and ease of use for researchers with diverse technical backgrounds. We demonstrate the utility of MassVision using an in-house generated dataset based on Desorption Electrospray Ionization (DESI) MSI of colorectal tissue, as well as public DESI and Matrix-Assisted Laser Desorption/Ionization (MALDI) MSI datasets. We also made our in-house dataset publicly-available to enable benchmarking, and facilitate comparative analysis to further advance the development of open-source tools for MSI data analysis.

## Design and Features

Figure 1 provides an overview of the end-to-end analytical pipeline in MassVision. The process begins with loading MSI data, after which users can create various visualizations to highlight the spatial distribution of different metabolomic properties. Corresponding histopathology images are then be co-localized with these visualizations to align the annotations with the spectra, guiding the labeling process. Spatial regions of interest may be selected to extract spectrum-label pairs, which may be exported as a curated dataset. Dataset curation can be performed independently for multiple slides, with all datasets seamlessly merged into a single dataset via feature alignment. The resulting multi-slide dataset can be preprocessed and utilized to train AI models for tissue characterization. Finally, trained models can be exported and deployed prospectively on new whole-slide MSI data.

**Figure 1:**
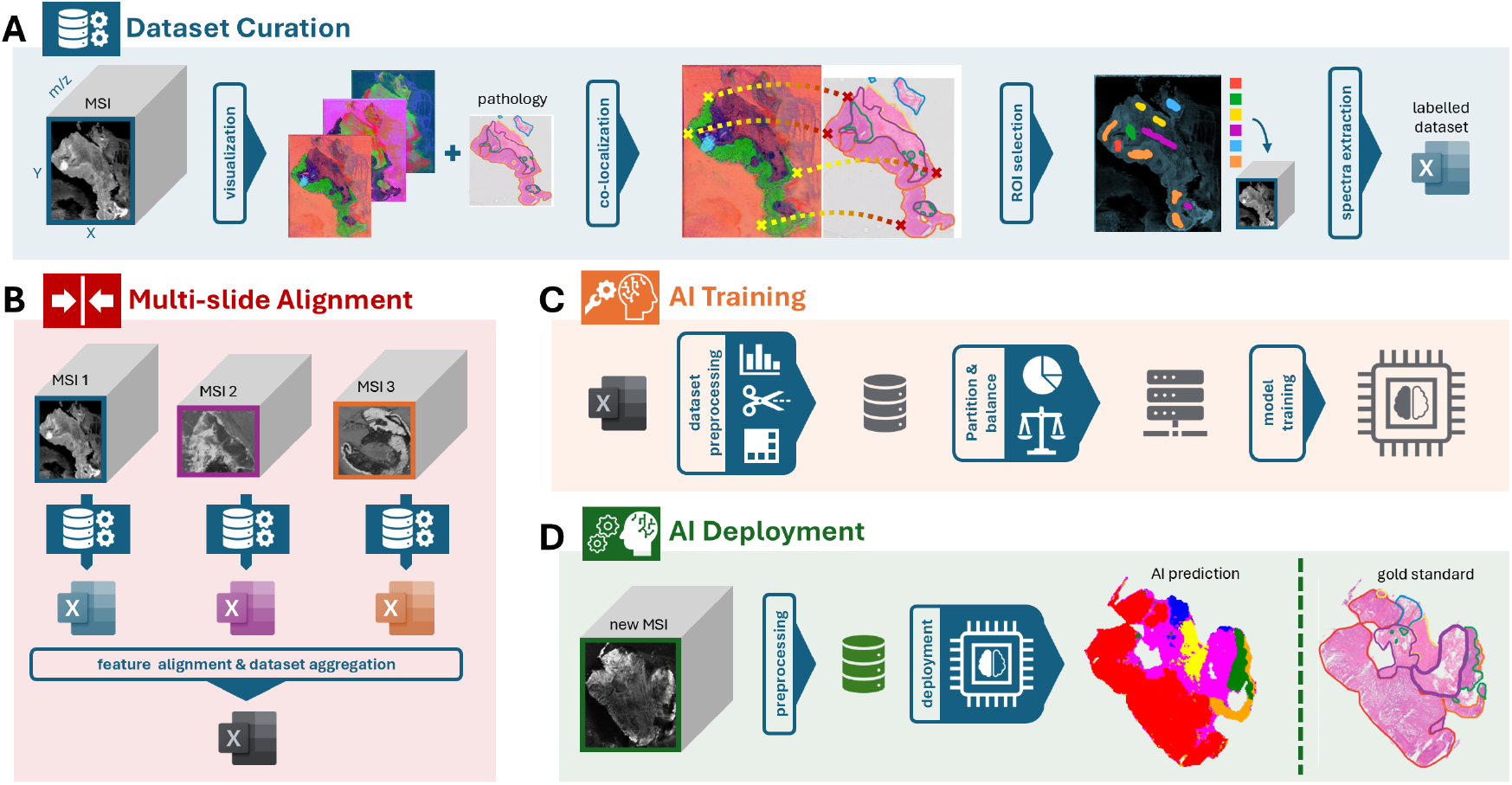
Overview of the end-to-end analytical pipeline in MassVision, demonstrating dataset curation (A), multi-slide alignment (B), AI model training (C), and deployment of AI models on whole-slide MSI (D).

### Implementation

MassVision is implemented as a module within the 3D Slicer platform, a widely-used opensource software suite for biomedical image analysis and visualization.^16,17^ The implementation of MassVision aligns with 3D Slicer’s standards, and the software is developed entirely in Python, a language widely adopted in scientific computing for its simplicity and versatility.^18^ The user interface is designed using Qt Designer, a widely used framework for creating modern and responsive graphical interfaces.^19^ By leveraging 3D Slicer’s well-documented APIs and development practices, MassVision ensures efficient integration and consistent performance within the platform. A view of the user interface of MassVision is demonstrated in supplementary Figure S1.

### Functionalities

MassVision organizes the functionalities required for end-to-end MSI analysis into multiple tabs, each tailored to streamline a specific stage of the workflow.

#### Data import

This tab allows users to load MSI data and associated files into MassVision. The current version supports modality-independent tabular CSV and hierarchical HDF5^20^ data formats, which accommodate rectilinear MSI data acquired from any source. Direct import of DESI text files generated by native instrument software is also available. Additionally, histopathology images may be loaded to guide the labeling of spectra within the dataset, or users may import previously saved projects, restoring all annotations and visualizations to resume analyses, conduct further experiments, or make modifications.

#### Visualization

Visualization refers to the abstract representation of rich MSI data as an image, providing interpretable insights into spatial patterns associated with key ions or metabolites. MassVision is equipped with various visualization approaches designed to facilitate data exploration and streamline dataset curation. Conventional methods include total ion current (TIC) map, that provides an overview of ion intensity across the sample; single-ion heatmaps that depict the spatial distribution of a selected ion; and multi-ion colormaps that overlay up to seven ions, each assigned a distinct color scale to reveal complex spatial relationships among metabolites. Additionally, spectrum plot for multiple pixels selected by the user is available.

To complement targeted methods that rely on prior knowledge of metabolites and relevant *m/z* values, MassVision introduces an untargeted approach for broader exploration of metabolomic distributions. This approach utilizes Principal Component Analysis (PCA) to reduce high-dimensional spectral data. The visualization is created by concatenating the first three principal components as the red, green, and blue channels of a color image, providing a comprehensive visual representation of the data. The unique innovation in MassVision lies in its ‘local metabolomic contrast’ visualization, where PCA is recalculated specifically for a spatial region on the slide defined by the user. By preserving the maximum data variance within the user-defined region during dimension reduction, the proposed local contrast visualization emphasizes dominant spectral features and reveals subtle yet significant patterns specific to the localized content.

For all untargeted visualizations, MassVision determines high-contributing ions identified through PCA loadings, providing deeper insights into the data. Additionally, users can apply clustering to these visualizations and retrieve the ions with strong correlation to each specific cluster, facilitating the ion identification.

#### Dataset curation

Generating well-annotated datasets with sample-label pairs is a fundamental step in preparing data for AI-driven analyses, ensuring robust training and validation of predictive models. For MSI, each data sample corresponds to a spectrum, where *m/z* values represent features, and the ion intensity of each specific *m/z* serve as feature value. Gold-standard labels are typically derived from a different modality, such as histopathology.

MassVision provides an integrated environment that enables users to perform multimodal image registration to co-localize and overlay pathology annotations with MSI visualizations to guide dataset curation (Figure 1A). The default registration method employs similarity transformation (shift, rotation, and scale without deformation) based on userdefined landmarks, while other validated registrations available within 3D Slicer can also be utilized.

After registration, MassVision integrates 3D Slicer’s Segment Editor module, allowing users to easily define regions of interest (ROIs) corresponding to each pathology label (Figure 1A). While simple paint and eraser tools are default options, automated segmentation methods within 3D Slicer can further streamline the ROI selection process. There are no limits on the number of class labels or data samples per each label, and each label group can be assigned a unique name and color for convenience. The spectra and corresponding labels of selected pixels are exported to a CSV file, with each row containing a spectrum, its label, and pixel location for spatial traceability of spectra throughout the analysis pipeline.

#### Multi-slide alignment

In AI-driven analyses, incorporating datasets from multiple patients is essential to account for variability and avoid mistaking patient-specific patterns for tissue-specific ones. In MSI, this process is challenging because each slide typically has a unique *m/z* list, preventing direct aggregation of slides. To address this, MassVision implements a feature alignment algorithm that unifies datasets before merging.

After importing curated datasets from different slides, MassVision extracts all *m/z* values and estimates their probability density function using kernel density estimation. The estimation parameters are tuned so each peak in the density plot represents a cluster of neighboring *m/z* values within a radius of 0.01 Da, forming a unified feature list. Spectra from each slide are then mapped to this reference list based on absolute ion distances. Once aligned, the datasets are merged into a single multi-slide dataset, which can be exported as a CSV file for further analysis.

#### Dataset preprocessing

MassVision is equipped with MSI-specific preprocessing algorithms designed to assist users in preparing data for downstream tasks. Users can normalize spectra to the total ion current (TIC) or the intensity of a reference ion. The *m/z* range can also be truncated to focus downstream analysis on a specific range of interest.

To reduce spectral noise, MassVision offers “pixel aggregation” that leverages the spatial location of pixels preserved in the dataset and combines spectra from neighboring pixels. Users can define the width of the neighboring window and select a statistical function (e.g., median, mean, minimum, or maximum) for aggregation. While aggregation improves the signal-to-noise ratio of spectra, it reduces the number of data samples in the dataset, as each sample now represents a summary of multiple pixels (spectra) rather than individual ones.

#### Model training

This tab facilitates dataset exploration, partitioning, balancing, and training of AI models. After loading a dataset, users can visualize the distribution of data samples in lower dimensions using a 2D PCA projection. Two scatter plots are provided in MassVision: one color-coded by slide to highlight inter-slide variability and another colorcoded by tissue labels to emphasize class-specific variability.

For model training, users can choose from three options: using the entire dataset for training, performing a random train-test split, or employing custom splits for fold-based or leave-one-out cross-validation. To mitigate biases from imbalanced datasets, users can balance the training data by down-sampling majority classes or up-sampling minority ones. Feature-based normalization is included in the pipeline, with normalization parameters derived from the training data. Available AI models in MassVision include PCA followed by Linear Discriminant Analysis (LDA), Random Forest (RF), Support Vector Machines (SVM), and Partial Least Squares Discriminant Analysis (PLS-DA). Model performance is summarized using confusion matrices and metrics such as accuracy, balanced accuracy, class-specific recall, and the Area Under the Receiver Operating Characteristic Curve (AUC) for both training and test sets. Additionally, users can save the trained AI models along with the *m/z* list (reference feature list) for prospective deployment.

#### Whole-slide MSI deployment

In addition to dataset curation and analysis, MassVision provides a platform for deploying trained models on whole slides, enabling the prospective use of MSI as a point-of-care histopathology tool—an essential step towards integrating MSI into clinical practice (Figure 1D). The Deployment tab supports the same data formats as the Import tab, requiring both the MSI slide and the model pipeline from the Training tab as inputs.

Users have the option to preprocess slides with normalization and pixel aggregation; *m/z* values are then aligned to the reference *m/z* list used for model training. The model is then applied to the slide, generating a visualization where each pixel is color-coded based on the predicted class of its spectrum. Users can also restrict deployment to specific regions of the slide using spatial masks. This feature is particularly useful for excluding background pixels and focusing analysis on tissue areas of interest.

### User requirements

A successful software platform extends beyond field-specific functionalities, meeting core user requirements such as availability, usability, flexibility, supportability, and maintainability. MassVision fulfills these requirements by being an open-source module within the 3D Slicer platform, ensuring cross-platform compatibility and public access to its source code. It features an intuitive interface designed for non-expert users, preserves spatial integrity throughout the analysis pipeline, and allows seamless saving and resuming of progress. Comprehensive documentation and modular design further enhance its accessibility, adaptability, and long-term usability. Detailed descriptions of these user requirements are provided in the Supplementary section.

## Application

In this section, we demonstrate the utility of MassVision in different use cases. We utilized an in-house DESI Time of Flight (TOF) MSI dataset from our previous study on colorectal cancer tissues.^21^ The dataset comprises DESI slides from cross-sections of 10 surgically resected samples. Each DESI data was lock-mass corrected and filtered to contain only the 2000 most abundant ions using the native instrument software. Each MSI slide was accompanied by corresponding a digital histopathology image, with precise spatial annotations of tissue subtypes of colorectal adenocarcinoma, benign mucosa, submucosa, smooth muscle, serosa, and inflammatory cells. The complete description of sample procurement and data acquisition can be found in the original publication.^21^ The DESI-MSI data and pathology images have been made publicly available specifically for MassVision,^22^ but the data can also be used separately for benchmarking and exploration with other tools.

### Data exploration using targeted and untargeted visualization

Exploratory data analysis is a crucial step in any study, providing a foundational understanding of the data prior to downstream analysis. In the context of MSI, data exploration can be greatly enhanced through informative visualizations that yield interpretable representations of the complex spatio-spectral information. An example of exploration via visualization using MassVision is depicted in Figure 2 for a representative DESI slide. Figure 2A displays the TIC visualization of the slide, with an inset in the lower left corner displaying the corresponding histopathology-annotated image for reference. TIC is the most basic visualization and offers the overall ion distribution and a coarse overview of tissue boundaries.

**Figure 2:**
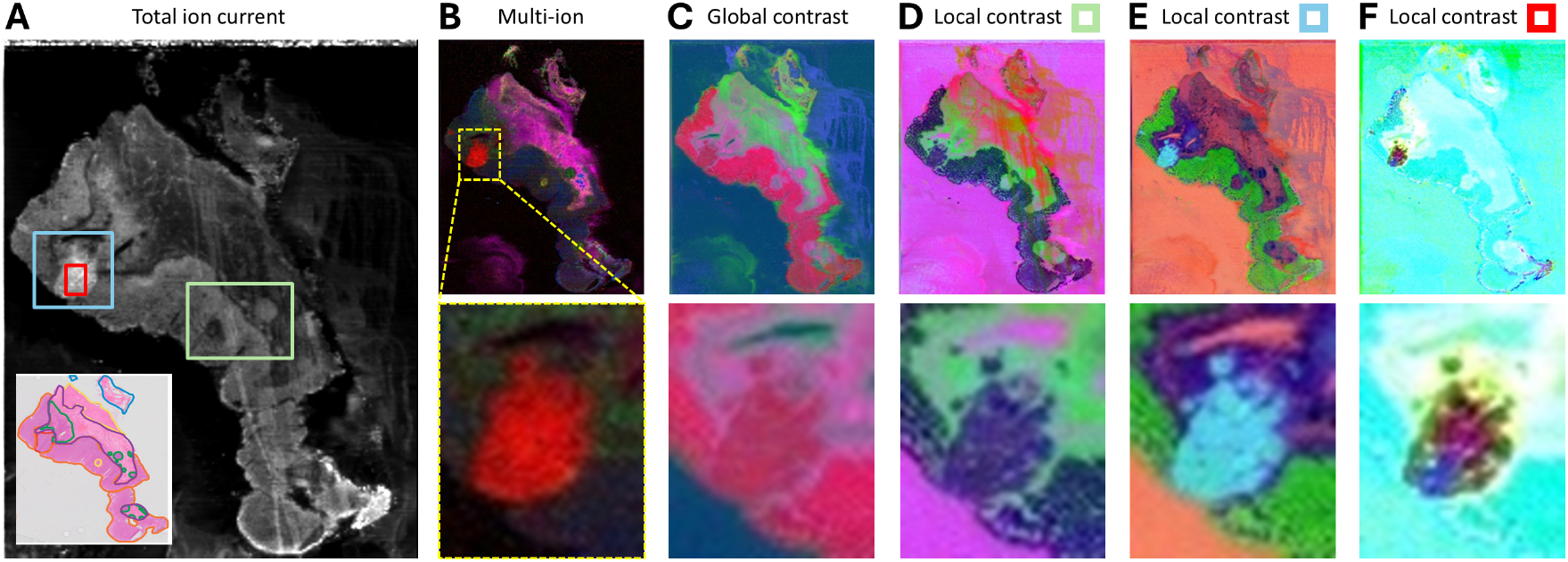
Demonstration of visualization methods in MassVision including TIC with an inset displaying the corresponding histopathology (A). multi-ion (B), global contrast (C), and local contrast visualizations optimized for green (D), blue (E), and red (F) boxes shown in TIC image, respectively. The bottom row for B-F represent a zoomed-in view of the region of interest.

A targeted multi-ion visualization of the slide is shown in Figure 2B, with the bottom row providing a zoomed-in view of the cancer region as determined by histopathology. Particularly, the red color scale represents the distribution of *m/z* 309.279 [FA(20 : 1) − H]^−^ (gondoic acid). This ion, found to exhibit high relative abundance in regions containing adenocarcinoma,^21^ serves as an example a well-correlated *m/z* value with histopathology. Such targeted visualizations are well-suited for spatial analysis of metabolites of interest that are already known to the analyst.

In the absence of prior knowledge about relevant metabolites, the proposed untargeted, region-specific visualizations, shown in Figure 2, offer a powerful tool for exploring MSI data. As demonstrated, while the global contrast visualization (Figure 2C)—optimized across all pixels—primarily accentuates metabolomic differences between tissue and background, it does not as distinctly differentiate adenocarcinoma from certain adjacent tissue types. By optimizing visualization parameters within a bounding box that encapsulates the cancer region, the metabolomic contrast specific to adenocarcinoma is significantly enhanced, as illustrated in Figure 2E.

In addition to inter-tissue contrast enhancement, the proposed untargeted visualization can reveal heterogeneity within a single tissue type. This is demonstrated in Figure 2F by defining a bounding box contained within the cancer region. While non-cancerous regions of the slide appear saturated and less informative, this visualization effectively highlights intratumoral heterogeneity. Understanding the metabolic diversity within cancerous tissue may reveal insights into tumor activity and mechanistic pathways that may be exploited as biomarkers or potential therapeutic targets.

### Pathology-validated identification of biologically-informative features

Spatial visualization of MSI data can effectively facilitate identification of specific ions that contribute to observed metabolomic diversity within tissues. Co-localizing these visualizations with histopathology annotations allows researchers to validate findings against goldstandard tissue typing, enhancing reliability and interpretability. This approach not only differentiates metabolic profiles across tissue types but also enables a detailed examination of variations within a single type. For example, in studying tumor heterogeneity, identifying ions associated with distinct sub-regions may provide insight into the molecular basis of intratumoral diversity, facilitating biomarker discovery and informing potential diagnostic and therapeutic strategies.^23^

To achieve this, we first co-localized the DESI slide with the histopathology image, using the global contrast visualization to better identify spatial landmarks within the tissue. Once aligned, we switched to the local contrast visualization to highlight the visible cancer heterogeneity identified in the previous section. Figure 3 presents a representative zoomed-in view of the cancer region on the DESI image alongside the co-localized pathology image. As shown, we identified three distinct metabolomic regions based on the color patterns in the visualization and plotted a representative spectrum of a pixel from each region, as illustrated in Figure 3D.

**Figure 3:**
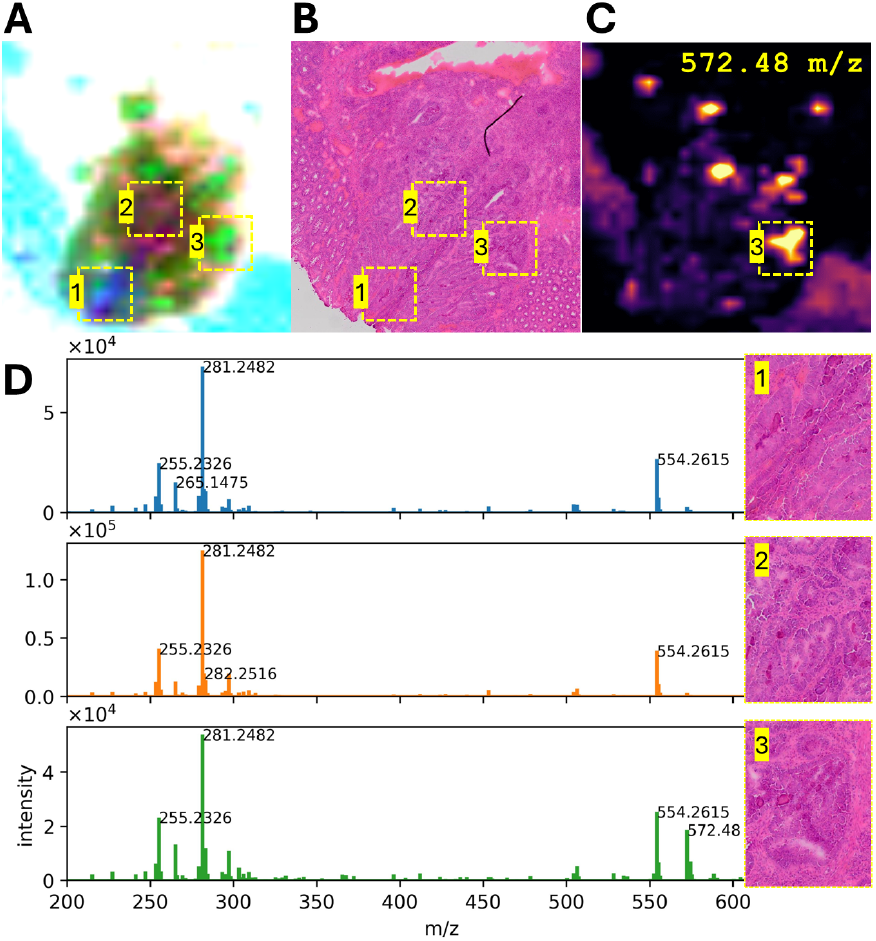
Pathology-validated feature identification in MassVision. A representative zoomed-in view of local contrast visualization (A), the co-localized pathology image (B), and the single-ion heatmap for *m/z* 572.48 (C), accompanied by three spectrum plots corresponding to representative pixels from the distinct regions highlighted in the visualization (D).

As seen, the spectra from regions 1 and 2 are quite similar, with the differences in metabolomic contrast not attributable to any specific ion. Pathological examination reveals that region 1 corresponds to more superficial tissue layers with larger glands, while region 2 represents deeper layers containing smaller glands. While the observed heterogeneity may result from metabolomic differences in tissue layering as previously reported in the literature,^24^ it may also be influenced by the imaging resolution. The average gland size in region 1 is 60 *μ*m while the DESI was collected in 100 *μ*m spatial resolution, contributing to a mixed spectral signal due to partial volume effects and hence different appearance in the visualization.

In contrast, as seen in last row of Figure 3D, the representative spectrum from region 3 displays a distinct peak at *m/z* 572.48, distinguishing it from other regions. Detailed pathological examination of this area reveals the presence of necrotic glands. The single-ion visualization of *m/z* 572.48 as shown in Figure 3C further supports this finding, as the ion’s abundance correlates closely with the distribution of necrosis observed in the histopathology image. Tata and colleagues also reported the correlation of *m/z* 572.48 [Cer(d34 : 1) + Cl]^−^ with necrotic cancer tissue present in human breast, which the group identified as a ceremide.^25^

It is important to emphasize that our initial histopathology-based assessment focused solely on delineating adenocarcinoma and other major anatomical sub-regions, rather than annotating region-specific heterogeneity. This example highlights the importance of MassVision’s untargeted, region-specific visualization tools in uncovering biologically relevant patterns that lack an initial annotation, but can later be correlated with underlying histopathological features (e.g., necrosis) through more detailed annotation.

### AI-based tissue-type classification

Tissue characterization using metabolomic signatures captured in MSI data plays a pivotal role in advancing our understanding of disease mechanisms and enhancing diagnostic precision. This process can be framed as a supervised classification task, where AI models are trained to learn feature patterns that differentiate between classes. Each data sample corresponds to a spectrum, and the class label represents the tissue type determined by a gold-standard method. Here, we demonstrate a use case for MassVision in the classification of tissue types in colorectal cross-sections.

After importing and visualizing each MSI dataset, the corresponding histopathology image with pathology annotations is co-localized to the MSI visualization using distinct tissue landmarks. MassVision links the registered pathology to the MSI data, enabling users to guide ROI selection effectively. A sample view of ROI selection in MassVision, along with the extracted regions for each tissue class, is shown in Figure 4. This process is repeated for all patients, with the resulting CSV datasets merged via feature alignment to create a single, multi-slide dataset.

**Figure 4:**
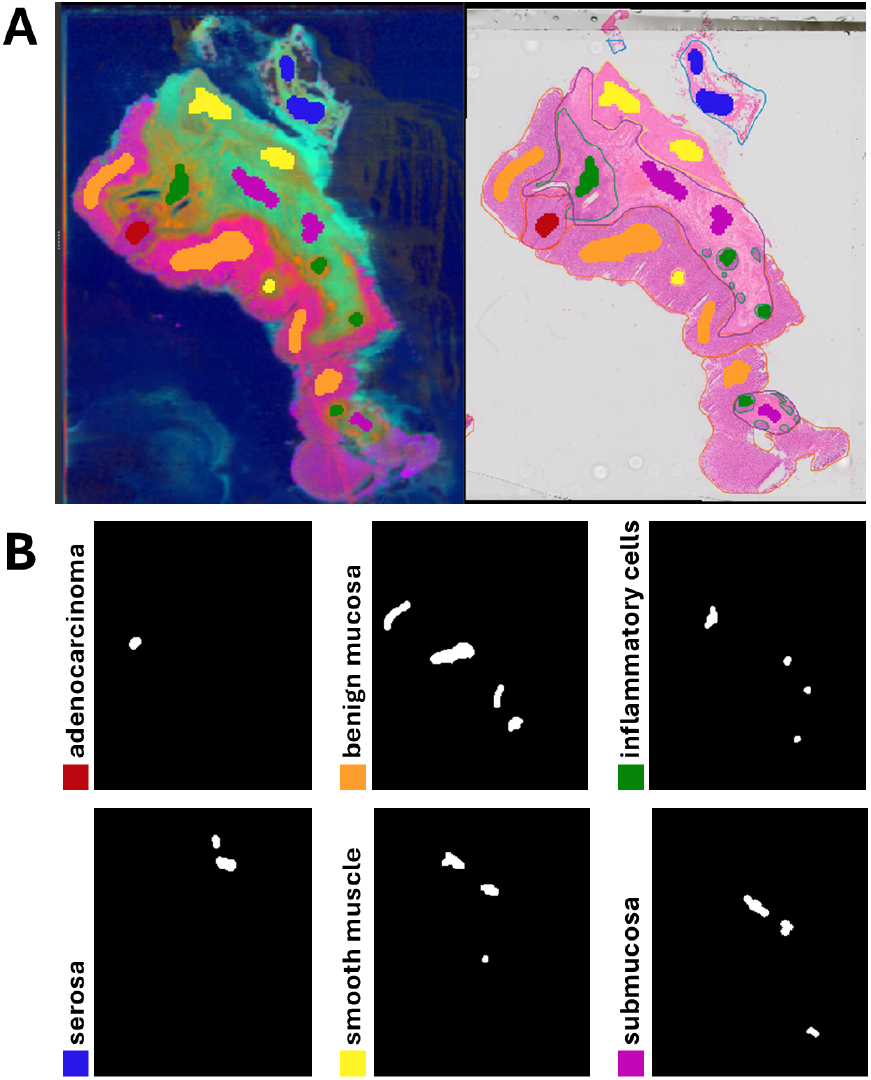
MassVision interface for ROI selection: (A) MSI visualization co-localized and linked with histopathology image annotated with six tissue subtypes of adenocarcinoma, benign mucosa, inflammatory cells, serosa, smooth muscle, and submucosa. (B) extracted regions of interest for each tissue subtype, color-coded according to the annotation.

For this study, the data were preprocessed using TIC normalization and pixel aggregation, where spectra were averaged within non-overlapping neighborhoods of 3×3 pixels. The dataset was split into 9 slides for training and 1 slide for testing. A PCA-LDA model was used for classification, and the effect of data balancing was evaluated on both the test dataset (quantitative validation) and the whole test slide (qualitative validation). Figure 5 presents the confusion matrices and the whole-slide deployment results for both the original and balanced models, alongside the annotated histopathology image of the test slide for reference.

**Figure 5:**
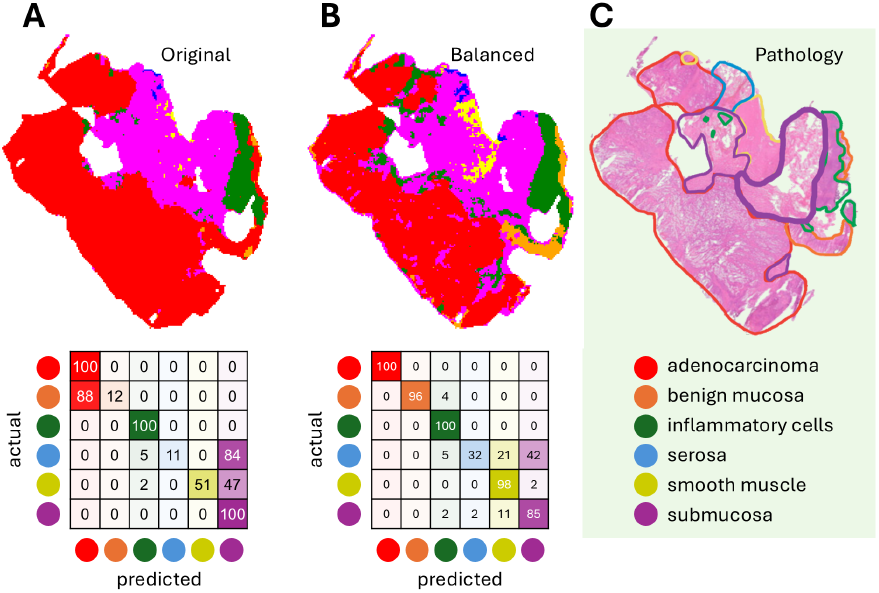
Qualitative and quantitative results highlighting the impact of data balancing: pixel classifications (top row) and confusion matrices (bottom row) for original (A) and balanced (B) models, with gold-standard histopathology reference (C). All confusion matrices are in percentage, and cells are color-coded according to the predicted tissue class.

The confusion matrix in Figure 5A highlights the tendency of original model to overpredict adenocarcinoma (for benign mucosa) and submucosa (for smooth muscle and serosa). These findings align with previous studies on the same dataset.^21^ The deployment results for the original model reveal that the slide is primarily classified into adenocarcinoma, inflammatory cells, and submucosa, with other subtypes largely missing. Examination of the training dataset indicates that the missing classes correspond to those with the fewest samples, highlighting a bias toward classes with more representatives. Balancing the data prior to training helps mitigate this issue, enabling better differentiation between classes.

In contrast, the confusion matrix of the balanced model in Figure 5B demonstrates a significant improvement in the accuracy of predicting benign mucosa. While the prediction of submucosa is slightly less optimal, there is a notable increase in the accuracy of serosa and smooth muscle classifications. A qualitative comparison of the balanced model wholeslide deployment results with the gold-standard histopathology further reveals substantial improvements in detecting minority classes, though the results are not a perfect match. These findings underscore the importance of data balancing in improving the performance and generalizability of classification models in MSI-based tissue characterization.

### Utility across MSI modalities

While the mentioned use cases primarily showcase analyses on our in-house DESI TOF MSI data, MassVision is modality-independent and compatible with any rectilinear MSI data that can be represented as a cubical spatio-spectral matrix. To demonstrate MassVision’s broad applicability, we conducted additional exploratory data analyses on three externally generated publicly available MSI datasets across various MSI platforms: (1) DESI Orbitrap MSI of human colorectal adenocarcinoma,^26^ (2) MALDI Fourier-Transform Ion Cyclotron Resonance (FT-ICR) MSI of human prostate cancer,^27^ and (3) MALDI FT-ICR MSI of mouse brain xenograft model of glioblastoma.^27^ The global contrast visualization of these data in MassVision is presented in Figure 6. All datasets were converted to the HDF5 format prior to analysis as instructed in the user guide.^28^ The results of exploratory analysis, presented in Supplementary Figures S2-S4, reveal how the application of MassVision tools recapitulates the colocalization of significant m/z features with cancerous and non-cancerous tissue that were reported in the original studies.

**Figure 6:**
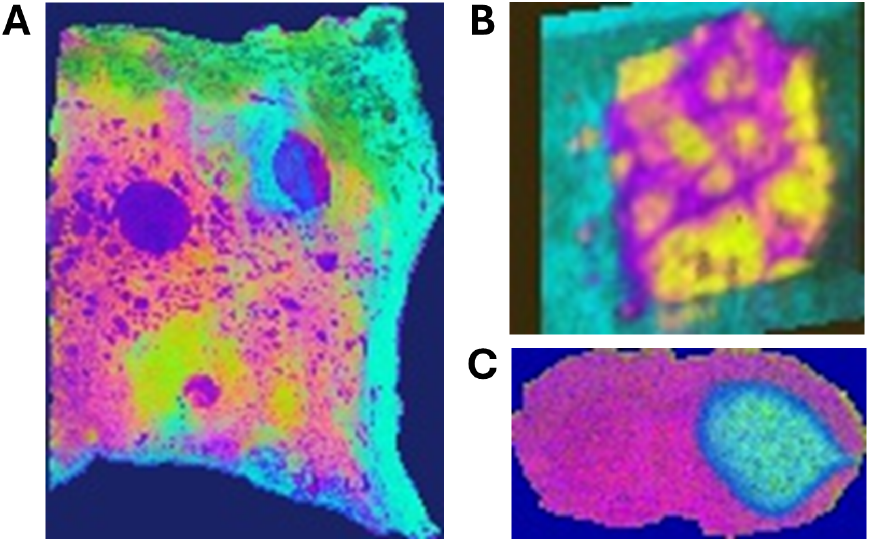
Global contrast visualization of publicly available data in MassVision: (A) MALDI of human prostate cancer, (B) DESI of human colorectal adenocarcinoma, and (C) MALDI of xenograft mouse brain.

## Discussion

Mass spectrometry imaging data provide a unique combination of spatial and spectral information, offering unprecedented insights into the molecular composition of biological samples. By enabling the visualization of molecular distributions across tissues, this technology has transformed applications in metabolomics, proteomics, and spatial biology. These advances not only support fundamental research into biological processes but also hold promise for clinical applications, such as tissue characterization and point-of-care histopathology. To fully harness this data-rich modality, robust software platforms are needed that integrate data curation, spatial information, and tissue classification into a single platform that is accessible to analysts without substantial data science expertise.

MassVision provides a comprehensive end-to-end solution for AI-based MSI analysis, addressing critical limitations in existing workflows. Unlike tools that focus on isolated tasks, MassVision integrates the entire workflow, eliminating the need for multiple platforms and the associated challenges with data transfer and file conversions. These intermediate steps can reduce pipeline productively and increase the risk of errors, particularly when handling large files. By streamlining these processes, MassVision ensures an efficient and reliable pipeline for MSI data analysis.

MassVision offers advanced features that enhance data exploration and analysis. Local contrast visualization highlights region-specific metabolomic variations, enabling the detection of subtle but significant patterns. Seamless pathology co-localization allows users to integrate histological annotations directly into MSI datasets for more precise analyses. Additional functionalities, such as guided labeling and multi-slide merging, support the creation of robust, multi-sample datasets while preserving spatial context. Together, these features bridge the gap between MSI data exploration and downstream analytical workflows, creating a unified analysis process.

3D Slicer offers an ideal platform for developing complex workflows and tools for MSI analysis. It features a modular architecture for developers and a user-friendly interface for non-programmers. Its built-in tools such as image registration and segmentation align well with MSI dataset curation requirements, reducing development overhead. Regular updates from a large and active community ensure continuous improvement and long-term sustainability. To the best of our knowledge, the only MSI module previously developed within Slicer is VIPRE.^29^ While VIPRE benefits from Slicer’s inherent strengths, it is private, limited to DESI analysis, features a suboptimal user interface, and lacks critical functionalities such as multi-slide merging and whole-slide deployment. MassVision builds on these strengths while addressing VIPRE’s limitations, introducing a versatile, user-friendly solution that bridges gaps in accessibility and functionality.

Table 1 summarizes the key features of leading tools for MSI analysis, highlighting their strengths and limitations alongside those of MassVision. In addition to its technical strengths, MassVision prioritizes usability and accessibility. Unlike High Definition Imaging (HDI)^12^ and MSiReader,^13^ which are commercial and not freely available, or Multi-MSIProcessor,^15^ which is restricted to specific operating systems, MassVision is built within the platform-independent open-source 3D Slicer ecosystem. This design ensures seamless integration into diverse research environments and eliminates the need for programming expertise, enabling researchers with varying skill levels to adopt the tool easily. By combining advanced capabilities with a user-friendly design, MassVision reduces barriers to adoption and fosters broader use of MSI in research and clinical applications.

**Table 1:** Comparison of key features across leading MSI analysis tools.

|  | HDI<sup>12</sup> | MSiReader<sup>13</sup> | MetaboAnalyst<sup>14</sup> | MMIPProcessor<sup>15</sup> | **MassVision (ours)** |
| --- | --- | --- | --- | --- | --- |
| Open source, free |  |  | ✓ | ✓ | ✓ |
| Platform independent |  |  | ✓ |  | ✓ |
| Spatial exploration | ✓ | ✓ |  | ✓ | ✓ |
| AI integration |  | ✓ | ✓ | ✓ | ✓ |
| Meets user requirements |  |  | ✓ |  | ✓ |
| Local contrast |  |  |  |  | ✓ |
| Pathology co-localization |  |  |  |  | ✓ |
| Guided data labeling |  |  |  |  | ✓ |
| Multi dataset merge |  |  |  |  | ✓ |
| Slide deployment |  |  |  |  | ✓ |

MassVision also demonstrates clear advantages in efficacy and throughput. Using the colorectal dataset as an example, the dataset curation workflow implemented with HDI in the original study^21^ required 6–8 hours per slide to produce 200–300 spectra. In contrast, applying the same workflow with MassVision reduced the time to 30–60 minutes while increasing the yield to 12,000–14,000 spectra per slide.

Additionally, MassVision preserves the spatial information and is highly scalable. For example, MetaboAnalyst,^14^ although a robust analytical platform, is limited in the size of datasets it can process due to its web-based design. It also lacks support for the integration of spatial information. In contrast, MassVision overcomes these limitations, efficiently handling large datasets while preserving spatial traceability throughout the entire processing pipeline. Overall, by combining cutting-edge functionality with user-centric design, MassVision accelerates the adoption of MSI in both research and clinical settings, bridging the gap between complex data and actionable insights.

## Conclusion

We introduced MassVision, a comprehensive, end-to-end solution for mass spectrometry imaging analysis that addresses critical challenges in workflow integration, data exploration, and analytical scalability. MassVision bridges the gap between molecular biology and computer science, enabling researchers without computing expertise to easily harness the power of MSI for their studies. Features such as local contrast visualization and pathology colocalization streamline dataset curation, enabling the efficient creation of high-quality, annotated datasets. Meanwhile, AI integration for model training and deployment extends the software’s utility to prospective studies, making MSI analysis practical for both research and clinical applications.

Moving forward, we envision MassVision evolving into an even more powerful tool through planned advancements and community-driven enhancements. Future developments include the integration of statistical analysis and deep learning algorithms, offering greater analytical flexibility to researchers. Additionally, as part of the 3D Slicer ecosystem, MassVision benefits from a dynamic user community whose feedback will shape future versions. This collaborative approach ensures the software remains adaptable, relevant, and aligned with the evolving needs of the MSI research community.

## Acknowledgement

A. Jamzad is supported by CIHR and NSERC grants. P. Mousavi is supported by a Canada Research Chair in Medical Informatics and Canada CIFAR AI Chair and the Vector Institute. G. Fichtinger is supported by a Canada Research Chair in Computer-Integrated Surgery. J. Rudan is supported by the Britton Smith Chair in Surgery. The authors thank Kyle Sunderland and Andras Lasso for their constructive comments and feedback during the implementation of MassVision.

## Supporting Information Available

## Availability Statement

The codebase for MassVision is publicly available on its GitHub repository ^1^. The documentations and detailed user guide are publicly available on MassVision website ^2^. The data used in this study will be publicly available upon acceptance of the paper.

## Supporting Information

Supplementary details on user requirement for the software; supplementary figures including user interface of MassVision; exploratory data analysis of three open-access data across different MSI modalities using MAssVision (PDF).

## TOC Graphic

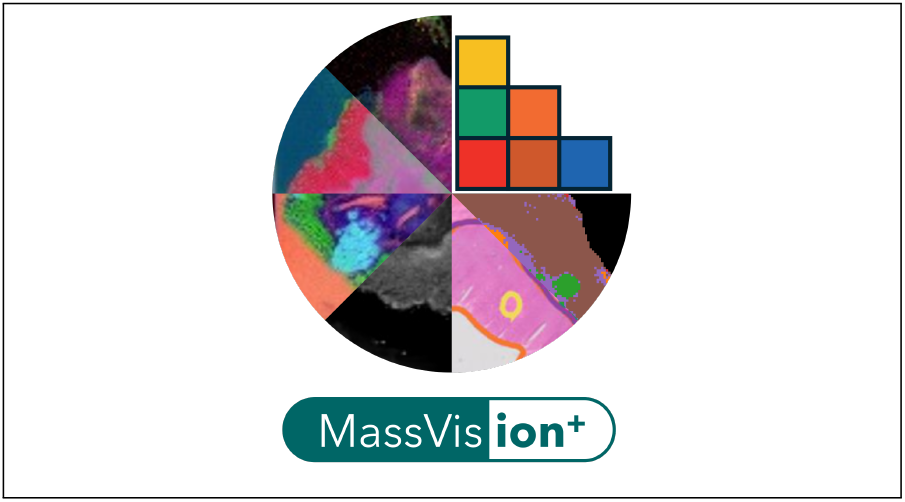

## Footnotes

1 https://github.com/jamzad/SlicerMassVision.git/

2 https://SlicerMassVision.readthedocs.io/

