## Supplementary for "MassVis*ion*: An open-source end-to-end platform for AI-driven mass spectrometry image analysis"

### Supplementary Information

#### User requirements

A successful software platform extends beyond specific functionalities, meeting core user requirements such as availability, usability, and flexibility to support a reliable and accessible data analysis pipeline for users with varied expertise levels. The following user requirements were identified and addressed in the development of MassVision.

**Availability:** As mentioned, MassVision is an open-source software developed as module within the 3D Slicer platform, ensuring straightforward access and easy installation for users. Its cross-platform compatibility allows for seamless installation on various operating systems, promoting broader accessibility and integration. Transparency is further supported through the public availability of the source code on GitHub, allowing users to review and understand the underlying algorithms and methodologies driving the software.

**Usability:** A primary development goal for MassVision was to bridge the gap between data scientists and bioanalytical chemists, allowing users with limited coding expertise to fully engage with the software tools. The interface is clean, intuitive, and user-friendly, with functionalities organized into clearly defined categories to make navigation straightforward for non-expert users. This user-centered design approach ensures that key features are easily accessible, facilitating efficient analysis workflows and minimizing the learning curve.

**Traceability:** Preserving the spatial integrity of the MSI data is essential for reproducible analysis. Unlike many existing platforms where spatial information is often lost during dataset curation, MassVision ensures that the pixel location of each spectrum within the slide is recorded and remains traceable across every step of the pipeline. Additionally, users can save the current state of their work, including all visualizations and ROIs, as a single project file, permitting the user to continue working on the same project at a later time. These features allow users to resume their work seamlessly, making it easy to review and modify ROIs as needed, and ensuring consistent and continuous data analysis.

**Documentation:** Comprehensive documentation is vital for ensuring that software tools are accessible, reliable, and effective for a wide range of users. For MassVision, detailed resources are provided, including a user manual, step-by-step tutorials, and practical examples, all available online. These materials guide users through each feature, allowing them to maximize the software's usability without requiring an extensive background in data science. Thus, MassVision tools are accessible for both new and experienced users.

**Flexibility:** Built within the 3D Slicer ecosystem, MassVision is designed with modularity and customizability in mind, enabling advanced developers to extend its core functionalities with Python coding. This flexible architecture enables ongoing adaptation and expansion to meet the evolving demands of research, making it a versatile foundation for future development.

**Maintainability:** Ensuring long-term usability and adaptability, MassVision is designed with robust maintainability practices. Version control through GitHub enables structured updates and tracking of changes, providing users with stable and accessible releases. Regular updates incorporate improvements and address user feedback, ensuring that the software remains relevant to evolving research needs. An active 3D Slicer community further supports collaborative development and input, helping sustain MassVision as a reliable and up-to-date tool for MSI data analysis.

### Supplementary Figures

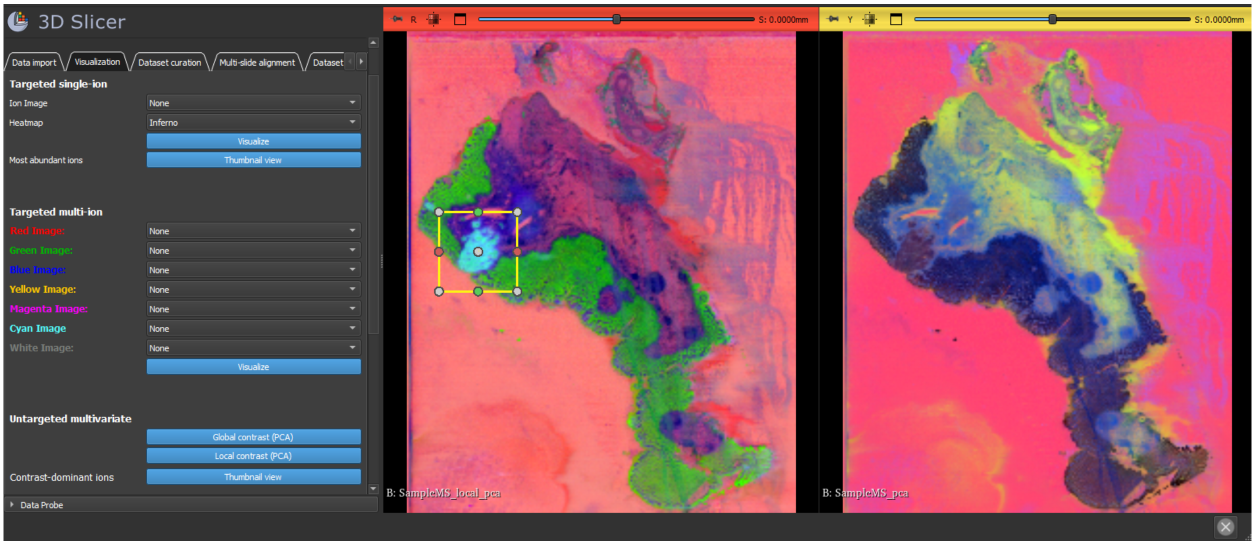


Figure S1: User interface of MassVision software demonstrating the visualization tab. The module panel on the left offers users access to implemented functionalities, while the view panels on the right are dedicated to visualizing analysis results.

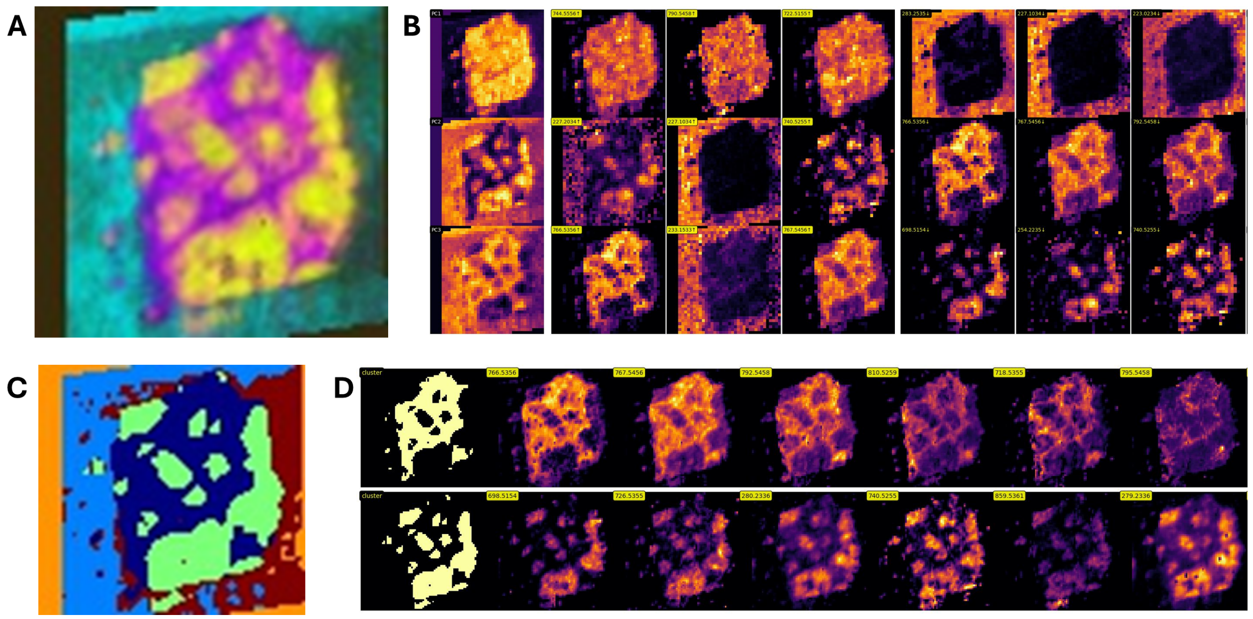


Figure S2: Exploratory data analysis of open-access DESI Orbitrap MSI data of human colorectal adenocarcinoma[1] using MassVision. The data matrix includes 5,694 pixels ($73 \times 78$) and 8,073 $m/z$ values per pixel. We demonstrated the following functionalities: (A) Global contrast visualization, (B) PC image, top positive loadings, and top negative loadings, (C) $K$-means clustering of global contrast visualization with $K$=5 clusters, and (D) clusters for normal (top row) and cancer (bottom row) tissues and corresponding ions with highest Pearson correlation. The observed increase in relative abundance of $m/z$ 279.2 and $m/z$ 766.5 within the tumor and normal clusters, respectively, have been previously reported.[2]

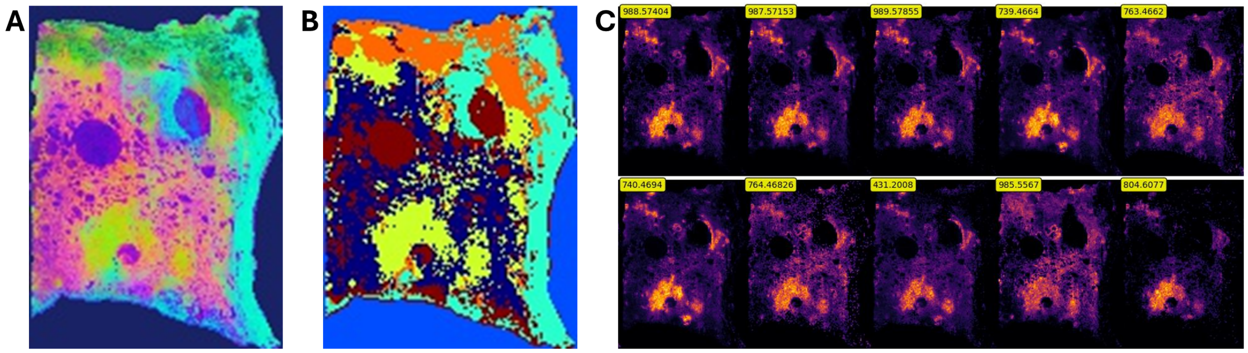


Figure S3: Exploratory data analysis of open-access MALDI FT-ICR MSI data of human prostate cancer[2] using MassVision. The data matrix includes 16,023 pixels ($147 \times 109$) and 61,343 $m/z$ values per pixel. We demonstrated the following functionalities: (A) Global contrast visualization, (B) $K$-means clustering of global contrast visualization with $K$=6 clusters, and (C) ions with highest Pearson correlation with cancer (yellow cluster in B). Among the reported ions, the correlation of $m/z$ 739.4664 (PA), $m/z$ 763.4662 (PA), , and $m/z$ 985.5567 PIP(P-42:6), with regions containing prostate cancer found by using MassVision, recapitulates the observation made in the original study.[2]

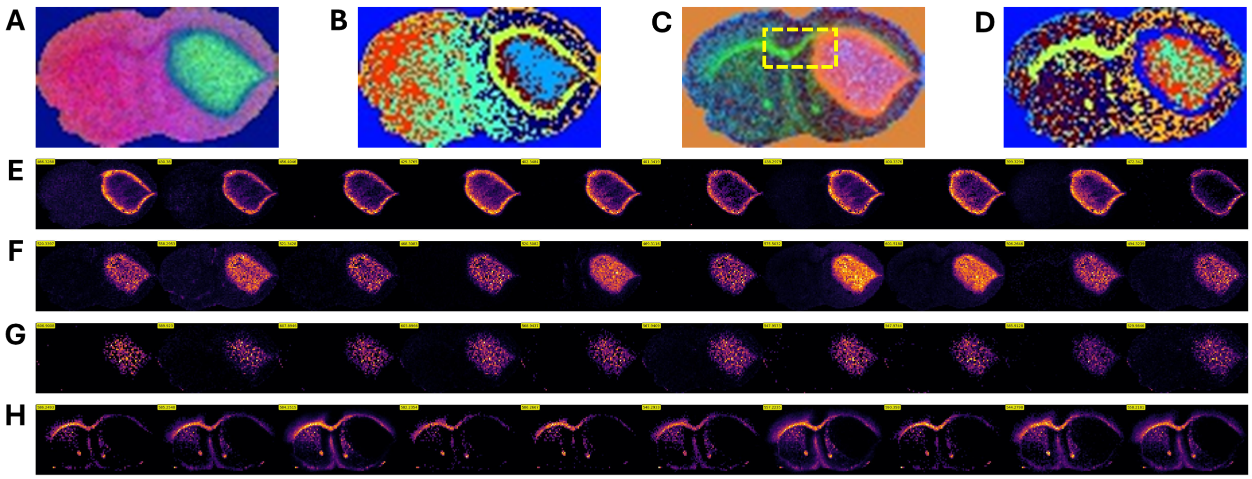


Figure S4: Exploratory data analysis of open-access MALDI FT-ICR MSI data of mouse brain xenograft model of glioblastoma[2] using MassVision. The data matrix includes 4,680 pixels ($52 \times 90$) and 21,241 $m/z$ values per pixel. We demonstrated the following functionalities: (A) Global contrast visualization, and (B) $K$-means clustering of global contrast visualization with $K$=8 clusters. The ions with high correlation to tumor rim and cancer clusters are presented in (E) and (F), respectively. (C) local contrast visualization optimized for dashed yellow region, and (D) $K$-means clustering of local contrast visualization with $K$=8 clusters. The local contrast better highlighted the heterogeneity of the tumor region (high correlated ions depicted in (G)) and revealed a new structure that was not visible in global contrast visualization (high correlated ions depicted in (H)). The significant ions found using the local and global contrast clusters in MassVision include $m/z$ 438.2979 (palmitoylcarnitine) which colocalized with the tumor rim region,[2] and $m/z$ 558.2953 (lysoPC(18:2)) and $m/z$ 529.9846 (ATP/dGDP) which were reported to correlate with tumor heterogeneity in the original study.[2]
